# Cell-based epitope screen identifies a novel B chain-derived Hybrid Insulin Peptide recognized by human and mouse autoreactive CD4 T cells

**DOI:** 10.64898/2026.09.16.752211

**Authors:** James E. DiLisio, Paul M. Zdinak, Nishtha Trivedi, Janet M Wenzlau, Anita C. Hohenstein, Joylynn Gallegos, Peter A. Gottlieb, Kathryn Haskins, Rocky L. Baker, Alok V. Joglekar

## Abstract

Type 1 diabetes (T1D) is mediated by autoreactive CD4 T cells that recognize pancreatic β-cell antigens MHC class II molecules. The B-chain of insulin is thought to be a primary autoantigen although native peptide sequences from the B-chain bind weakly to disease-associated MHC risk alleles. Hybrid Insulin Peptides (HIPs), consist of proinsulin fragments fused with peptides from other β-cell proteins, and may endow B-chain peptides with C-terminal acid residues that allow strong binding to MHC risk alleles and subsequent stimulation of autoreactive T cells. Here we applied a high-throughput cell-based screening platform with Signaling and Antigen-presenting Bifunctional Receptors (SABRs) to interrogate a theoretical library of insulin B-chain HIPs against clonally expanded CD4 T cell receptors from NOD mouse islets. We identified a previously undescribed HIP containing a fragment of insulin B-chain combined with a calreticulin-derived peptide. I-A^g^^7^ tetramers loaded with the InsB/Calr HIP bind to T cell clones as well as islet-infiltrating CD4 T cells and CD4 T cells from a recent-onset HLA-DQ2 T1D donor reacted exclusively to the InsB/Calr HIP. Collectively, these results demonstrate that insulin B-chain-derived peptides can undergo post-translational modification through HIP formation to generate neoepitopes with increased antigenicity, and underscore the utility of cell-based screening for identifying disease-relevant neoepitopes in T1D.

## Introduction

Type 1 diabetes (T1D) is an autoimmune disease characterized by T cell mediated destruction of insulin-producing pancreatic β-cells. Autoreactive CD4 T cells recognizing β-cell peptides presented by MHC class II molecules are thought to play a central role in disease pathogenesis by orchestrating inflammatory responses within pancreatic islets and providing help to cytotoxic CD8 T cells. Among the β-cell antigens implicated in T1D, insulin is a well-characterized target of autoreactive T cell responses, with multiple studies identifying insulin-derived peptides as ligands for pathogenic CD4 T cells in NOD mice^1–3^ and humans with T1D^4,5^.

Within the insulin molecule, the B-chain has emerged as a dominant region for T cell recognition^2,6^. Peptides spanning residues B:9-23 represent one of the most well characterized epitopes recognized by diabetogenic CD4 T cells in the NOD model and in humans. Structural and functional studies have shown that this peptide binds the disease-associated MHC class II molecule I-A^g^^7^ in alternative registers capable of activating autoreactive T cells that escape thymic deletion ^7–11^. Importantly, T cells recognizing the same B:9-23 peptide presented in high-risk HLA class II alleles are also detectable in humans with T1D^12–14^. However, native insulin B-chain peptides bind disease-associated MHC/HLA class II molecules with relatively low affinity^7,15,16^, suggesting that additional modification may be required to enhance their presentation to autoreactive T cells.

More recently, it has become clear that T cells targeting post-translational modifications (PTMs) may be key drivers of multiple autoimmune diseases^17–20^. Hybrid insulin peptides (HIPs) are a class of post-translationally modified epitopes that arise through covalent fusion of a fragment of the insulin protein with peptides derived from other β-cell proteins, generating neoepitope sequences that are not encoded in the genome. HIPs were first identified as targets of diabetogenic CD4 T cells in the NOD mouse, with corresponding HIP-reactive CD4 T cell responses also detected in individuals with T1D^21^. Subsequent work demonstrated that HIP-reactive T cells are present in humans with T1D^22–27^ and that hybrid epitopes can be enzymatically generated through fusion of proinsulin fragments with peptides derived from β-cell secretory granule proteins^28^. These findings identify HIPs as an important class of β-cell-derived neoepitopes in T1D^29^ and suggest that insulin fragments, including B-chain sequences, can participate in HIP formation to generate antigenic ligands recognized by autoreactive T cells, consistent with our previous studies ^11,30^.

Despite these discoveries, systematic identification of HIP epitopes and the disease-relevant TCRs that recognize such neoantigens remains challenging because the number of potential peptide fusion combinations is extremely large. Traditional synthetic peptide screening approaches are limited in throughput and cannot efficiently interrogate thousands of candidate epitopes against disease-relevant TCRs^19,31^. To address this challenge, new high-throughput antigen discovery technologies have been developed^31^. We have previously described a cell-based epitope discovery method that relies on chimeric receptors called Signaling and Antigen-presenting Bifunctional Receptors (SABRs). SABRs fuse a covalently linked peptide-MHC complex with an intracellular signaling domain, allowing presentation of a defined epitope to T cells and induction of a readable signal upon its recognition. SABR libraries provide a powerful platform for antigen discovery in mice and humans^32–34^. by enabling rapid functional screening of large defined peptide libraries presented in disease-relevant MHC molecules against TCRs of interest. Using SABR libraries presenting theoretical HIPs formed from the C-peptide of proinsulin, we previously identified novel InsC-Scg1 HIPs that were recognized by islet-infiltrating CD4 T cells in NOD mice^35^.

In the present study, we applied SABR screening to identify HIPs derived from the insulin B-chain that are recognized by clonally expanded CD4 T cells isolated from NOD islets. Using a theoretical B-chain HIP library, we identified a previously undescribed insulin B-chain HIP capable of activating multiple InsB:9-23-reactive TCRs. We further characterized the residues required for antigenicity, identified B-chain HIP-specific CD4 T cells within NOD islets, and demonstrated that human CD4 T cell clones isolated from a recent-onset T1D donor respond to this B-chain HIP neoepitope but not to the corresponding native B-chain sequence.

## Results

### Clonally expanded, insulin B-chain-reactive TCRs from islets of NOD mice

To define the antigen specificity of CD4 T cells infiltrating pancreatic islets during autoimmune diabetes, we reanalyzed single-cell RNA and paired TCRαβ sequencing on CD4 T cells isolated from pancreatic islets of prediabetic NOD mice^33^. After removing naïve clusters that had few expanded clones, dimensional reduction of the transcriptomic dataset revealed multiple transcriptionally distinct clusters representing phenotypically diverse CD4 T cell populations within the islet microenvironment (**Fig. S1A,B**). These clusters included populations expressing transcriptional signatures consistent with stem-like (0), effector (1,3,4), regulatory (2), and proliferative (5) T cell states. Overlay of expanded TCR clonotypes onto the transcriptional landscape revealed clonal populations occupying multiple T cells states (**Fig. 1A**), indicating antigen-driven proliferation and differentiation of T cells within the target tissue. Among the expanded clonotypes were three TCRs previously shown to recognize InsB:9-23 peptide using SABR libraries^33^, designated TCR11, TCR24, and TCR30.

**Figure 1.**
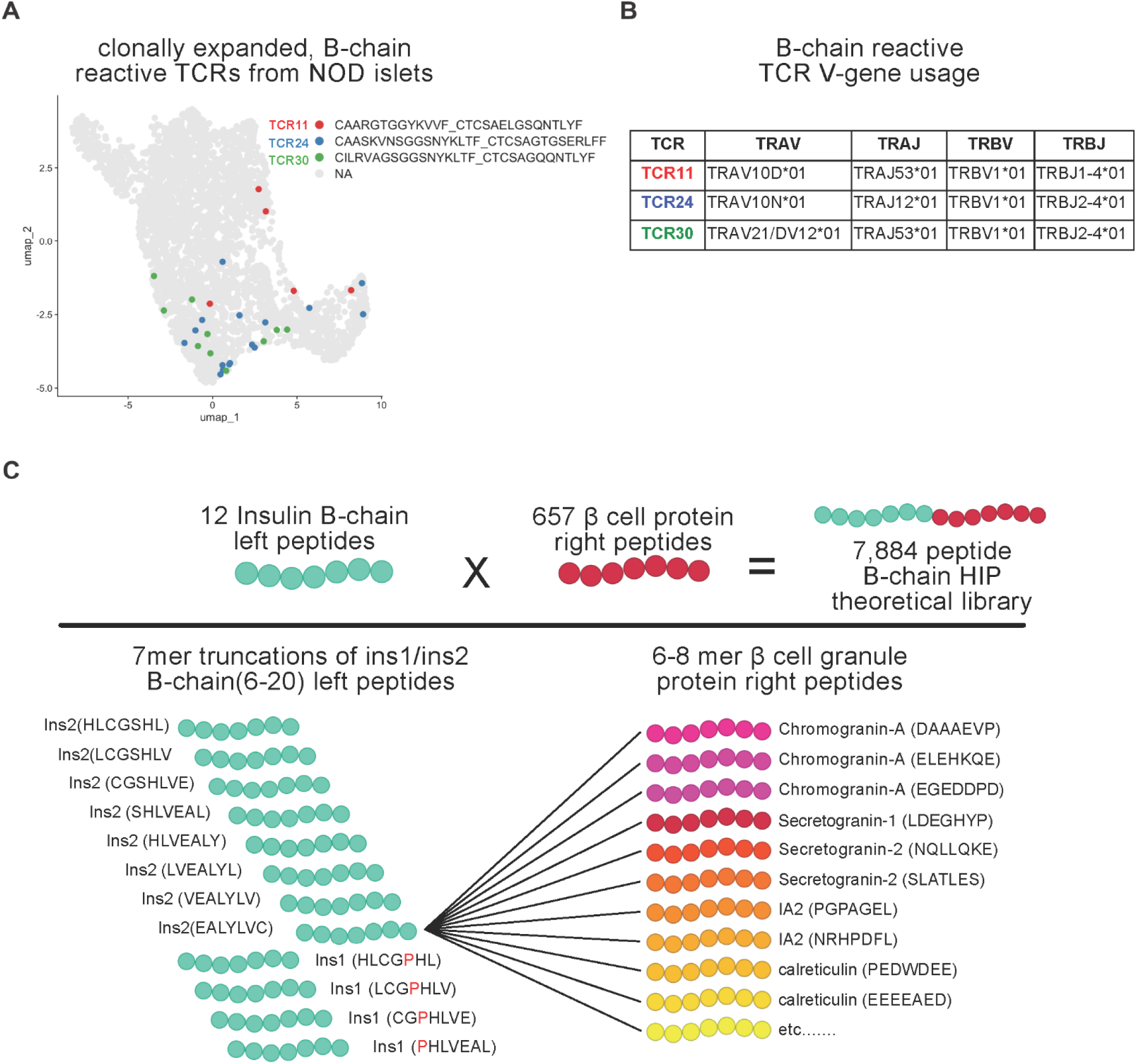
Clonally expanded, insulin B-chain-reactive TCRs and theoretical B-chain hybrid insulin peptide library. (A) uMAP projection of single-cell RNA sequencing data from CD4 T cells isolated from pancreatic islets of prediabetic NOD mice (8–12 weeks of age). Clonally expanded insulin B-chain-reactive TCR clonotypes (TCR11, TCR24, and TCR30) identified by paired (TCRα_TCRβ) CDR3 sequences are highlighted in red (TCR11), blue (TCR24), and green (TCR30). (B) TCR gene segment usage for expanded clonotypes TCR11, TCR24, and TCR30. (C) Schematic illustrating construction of the theoretical insulin B-chain hybrid peptide (HIP) library. Seven-amino acid truncations spanning insulin B-chain residues 6-20 derived from Ins1 and Ins2 sequences were paired with 6-8 amino acid fragments derived from β-cell secretory granule proteins including insulin itself there were identified by mass-spec of islets^44^, beta cells^45^, and eluted from the intra-islet immunopeptidome^37^. Combinatorial pairing of each B-chain left peptide with all 657 β-cell granule protein right peptides generated 7,884 theoretical, candidate B-chain hybrid insulin peptides.

Analysis of TCR gene usage revealed notable similarities among the clones. All three TCRs utilized TRBV1, although their TRAV and TRAJ gene segments differed (**Fig. 1B**). TCR11 and TCR30 shared identical TRAJ gene usage and displayed related CDR3α motifs (**Fig. S1C,D**), possibly suggesting structural convergence in antigen recognition. Such convergent TCR features have previously been observed among insulin-reactive T cells and may reflect selective pressure imposed by presentation of specific insulin-derived epitopes^36^. Given the central role of insulin B-chain epitopes in autoimmune recognition, we hypothesized that hybrid peptides derived from insulin B-chain fragments might serve as the endogenous ligands for these expanded TCRs.

### Construction of a putative B chain HIP SABR library

To systematically interrogate this hypothesis, we constructed a theoretical B-chain HIP library. The N-terminal component of the library consisted of 7mer truncations spanning insulin B-chain residues 6-20, derived from both insulin-1 and insulin-2 sequences. This region encompasses the well-characterized B:9-23 epitope and adjacent residues implicated in T cell recognition. The C-terminal component of the HIP consisted of 7mer peptides derived from proteins enriched in β-cell secretory granules^37^ including proinsulin, chromogranin A, secretogranin-1, secretogranin-2, IA-2, calreticulin, and other proteins found in β-cells. Combinatorial pairing of insulin fragments with β-cell peptide fragments generated a library of 7,884 candidate peptides (**Fig. 1C**). Together, this design generated a comprehensive library of putative B-chain HIPs enabling systematic interrogation of potential neoantigens recognized by islet-derived CD4 T cells. The 7,887 epitope B-chain HIP library, including three control peptides, was cloned in the I-A^g^^7^ SABR backbone using a pooled oligonucleotide cloning strategy at >100x coverage^34^ and expressed in NFAT-GFP-Jurkat cells (**Fig 1D**).

### InsB:9-23-reactive TCRs cross-react with a novel HIP composed of insulin B-chain and a fragment of calreticulin

We screened Jurkat cells expressing TCR11, TCR24, or TCR30 against the B-chain HIP SABR library. To screen the cells, TCR-expressing Jurkat cells and SABR-expressing NFAT-GFP-Jurkat cells were cultured for 18 hours at 1:1 ratio. Following co-incubation, GFP+CD69+ NFAT-GFP-Jurkat cells were FACS sorted and genomic DNA was extracted. Using our published protocol^33^, the epitope-encoding portion of the SABR backbone was amplified and sequenced.

Screens for each TCR were performed in triplicates, with the unsorted library acting as the baseline control. An Enrichment Score was calculated for each epitope within each TCR screen (**Fig 2B, S2A**). Analysis of enriched sequences following sorting of GFP-positive cells revealed several B-chain HIPs that suggested cross-reactivity with the TCRs (**Fig. 2B**). These HIPs were confirmed as containing the N-terminal insulin B-chain residues EALYLVC and C-terminal fragments from calreticulin (PEDWDEE), Secretogranin 1 (SEEEPEY), and carboxypeptidase E (CNDDDSSF) (**Fig 2B**). Using single SABRs corresponding to these epitopes, the reactivity of the individual TCRs towards those epitopes was validated (**Fig 2C**). These results show that InsB:9-23-reactive TCRs show cross reactivity towards putative B-chain HIPs. We chose the InsB/Calr peptide (EALYLVC/PEDWDEE) for further analysis as it stimulated both TCR11 and TCR30 expressing cells. To test whether reactivity to InsB/Calr HIP reactivity was found in TCRs beyond the two that were screened, several TCR analogs previously identified through TCR-similarity metrics^33^ were also tested for their ability to recognize this peptide. These TCR analogs were identified as putatively reactive to InsB:9-23 through the computational TCR prediction algorithms, tcrdist3^38^ and GLIPH2^39^, and several of them were validated for their reactivity to InsB:9-23 peptide. A total of 5/13 analogs showed reactivity to the InsB/Calr HIP. Interestingly, 4/5 analogs as well as TCR11 and TCR30 showed higher reactivity to the InsB/Calr HIP than to InsB:9-23 (**Fig 2D**). These data suggest that InsB/Calr HIP is an islet neoepitope that is recognized by the insulin B-chain-reactive CD4 T cell repertoire.

**Figure 2.**
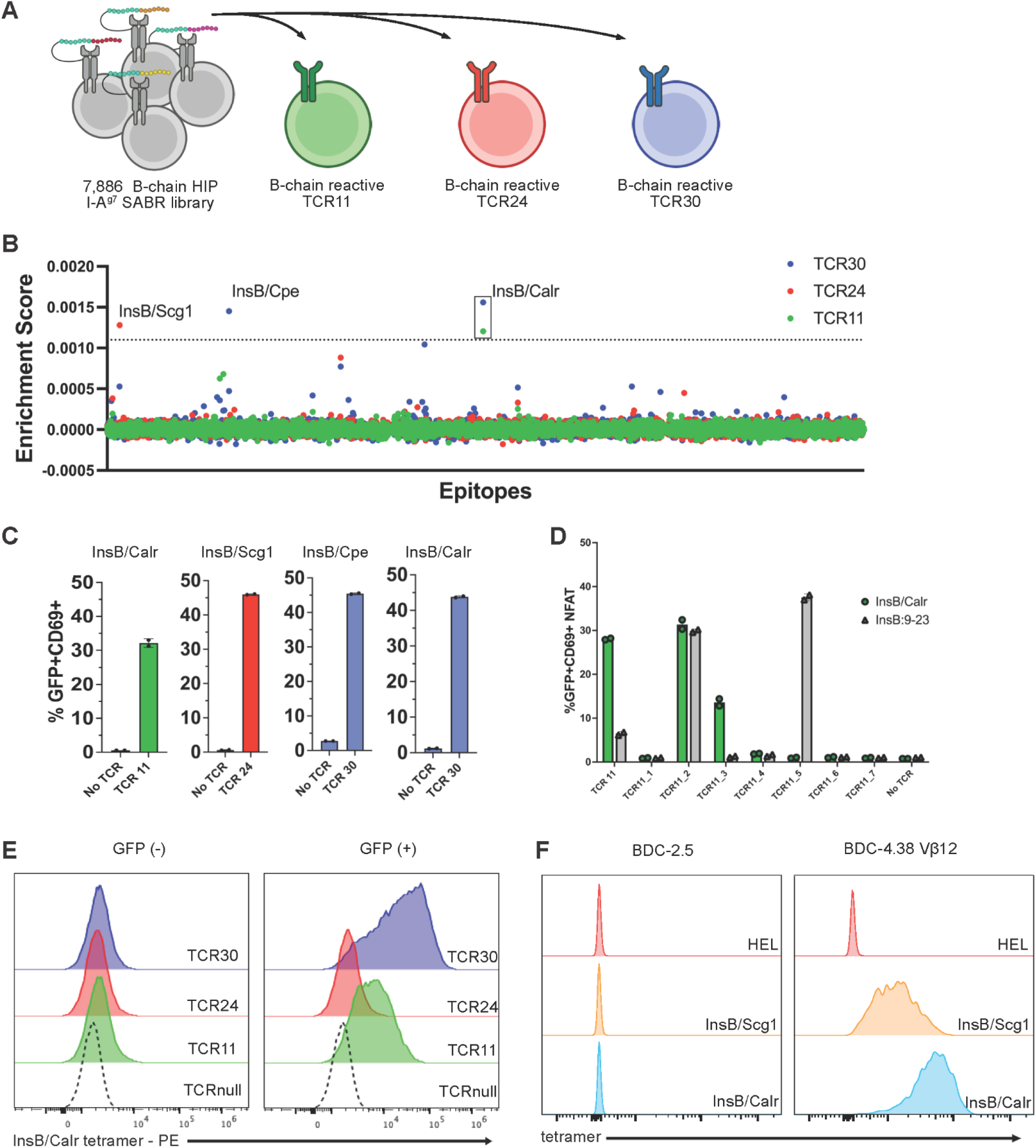
SABR screening identifies a hybrid insulin peptide composed of insulin B-chain and a fragment of calreticulin. (A) Experimental workflow for SABR-based screening. A 7,884 B-chain HIP library encoded within I-A^g^^7^ SABR constructs were expressed in NFAT-GFP reporter Jurkat cells and co-cultured with individual TCR-expressing Jurkat cells for TCR11, TCR24, and TCR30. Engagement of peptide-MHC complexes by cognate TCRs triggered intracellular signaling resulting in GFP reporter activation which is normalized by an enrichment score relative to the pre-co-culture abundance. (B) Enrichment analysis of B-chain HIPs following SABR screening. Candidate peptides enriched among GFP+ reporter cells are plotted. The InsB/Calr hybrid peptide (EALYLVC/PEDWDEE) was selectively enriched in screens performed with TCR11 (green dots) and TCR30 (blue dots) but not TCR24 (red dots). (C) Single SABR (InsB/Calr, InsB/Scg1, InsB/Cpe) assays with TCR11, TCR24, or TCR30. (D) InsB/Calr or native InsB9-23 single SABR assays with TCR11 and putative TCR analogs 1-7. (E) I-A^g^^7^ tetramer staining of Jurkat cells expressing TCR11, TCR24, and TCR30 in GFP-(no construct) and GFP+ (TCR construct) populations. (F) HEL, InsB/Scg1, or InsB/Calr I-A^g^^7^ tetramer staining of BDC-2.5 or BDC-4.38 Vβ12 CD4 T cell clones.

To further confirm antigen specificity outside of the SABR system, an I-A^g^^7^ tetramer loaded with the InsB/Calr HIP was generated by the NIH Tetramer Core Facility (insB/Calr HIP tetramer). Based on prior studies demonstrating increased TCR reactivity to B-chain HIPs containing the N-terminal valine^30^, the tetramer was synthesized using the sequence (V)EALYLVC/PEDWDEE. This tetramer moderately stained TCR11-expressing cells and robustly stained TCR30-expressing cells, while showing no detectable binding to TCR24 above background (**Fig 2E**, gating in **Fig 3A**). Furthermore, the insB/Calr HIP tetramer stained BDC-4.38 Vβ12 (**Fig. 2F**), an insulin-reactive, diabetogenic CD4 T cell clone from the BDC panel ^40^. This tetramer did not stain the BDC-2.5 CD4 T cell clone, which is reactive to a C-peptide/ChgA HIP^21^, and of note, the BDC-4.38 Vβ12 T cell clone did not stain with either the insp8G or the insp8E tetramer (**Fig. S4A**). Together, these results confirm that the InsB/Calr HIP is a peptide capable of activating multiple B-chain-reactive TCRs derived from the islet infiltrate and can be specifically detected using peptide-MHC tetramers.

**Figure 3.**
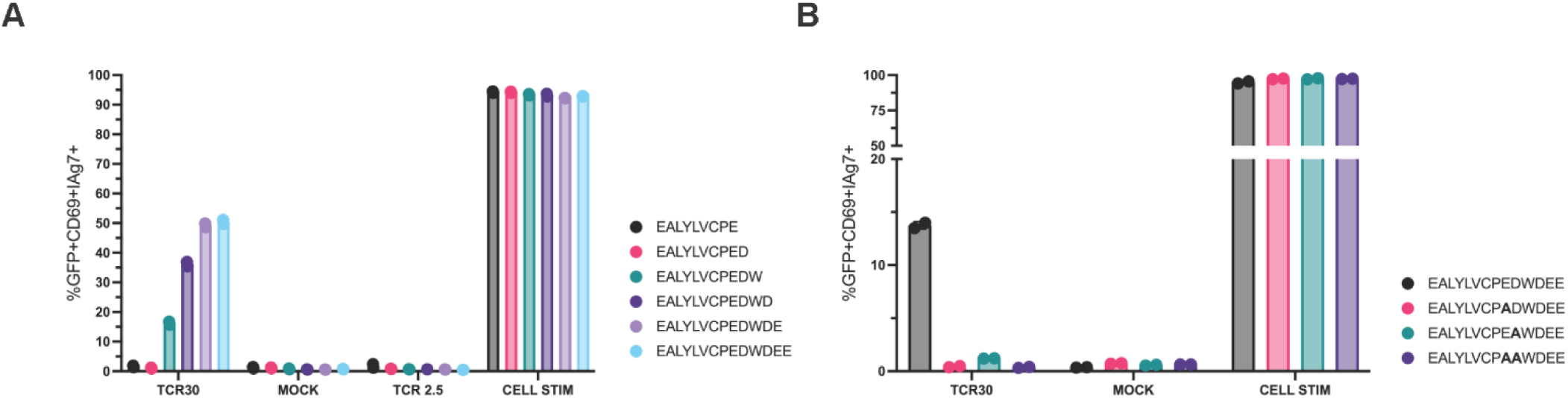
Acidic residues within the calreticulin-derived segment determine antigenicity of the InsB/Calr hybrid peptide. (A) Activation (NFAT-GFP+CD69+) of SABRs baring N-terminal truncation variants of the InsB/Calr HIP incubated with TCR30. (B) Activation (NFAT-GFP+CD69+) of SABRs baring alanine substitution at potential p9 pocket residues of the InsB/Calr HIP incubated with TCR30.

### Acidic residues within the calreticulin-derived segment determine antigenicity of the InsB/Calr HIP

To determine which residues within the calreticulin-derived segment contribute to antigen recognition, we generated truncation and alanine substitution variants of the InsB/Calr peptide. Single SABR assays demonstrated that progressive truncation of residues within the calreticulin-derived segment markedly reduced TCR 30 activation (**Fig. 3A**). These results indicated that C-terminal residues, including the tryptophan (W), contributed by calreticulin are required for optimal antigenicity through enhanced binding of MHC or stimulation of the TCR.

Acidic residues are favored at the p9 anchor position of the I-A^g^^7^ peptide-binding groove, where they stabilize peptide-MHC interactions^41^. Alanine substitutions at residues predicted to occupy the p8 and p9 registers of the peptide-MHC complex also significantly reduced activation of TCR30 (**Fig. 3B**). Taken together, these results highlight the importance of the hybrid peptide junction in determining antigenicity and suggest that C-terminal addition of the acidic D/E residues to B-chain fragments may endow the resulting HIP with increased MHC binding and subsequent recognition by TCRs that otherwise weakly recognize native B-chain sequences.

### InsB/Calr-specific CD4 T cells are present within pancreatic islets of NOD mice

To determine whether T cells recognizing the InsB/Calr peptide exist *in vivo*, we performed dual tetramer staining using PE and APC-labeled I-A^g^^7^ tetramers loaded with the InsB/Calr peptide on islet-infiltrating T cells. Flow cytometric analysis of CD4 T cells isolated from NOD islets revealed a distinct population of PE+APC+ dual-tetramer staining cells (**Fig. 4A** gating in **S3B**). InsB/Calr tetramer+ T cells were detected among islet-infiltrating CD4⁺ T cells at frequencies comparable to other insulin-specific populations that stained with the insp8E and insp8G tetramers (**Fig. 4B**). In most NOD mice, InsB/Calr HIP tetramer+ cells stained a population distinct from insp8E- and insp8G-reactive T cells. However, in some animals there was a subset of T cells that stained with both the InsB/Calr HIP and p8G tetramers (**Fig. S5A**). These observations suggest that some TCRs are capable of recognizing both native insulin B-chain peptides and InsB/Calr HIP derived-epitopes, whereas some TCRs only recognize the HIP epitope.

**Figure 4.**
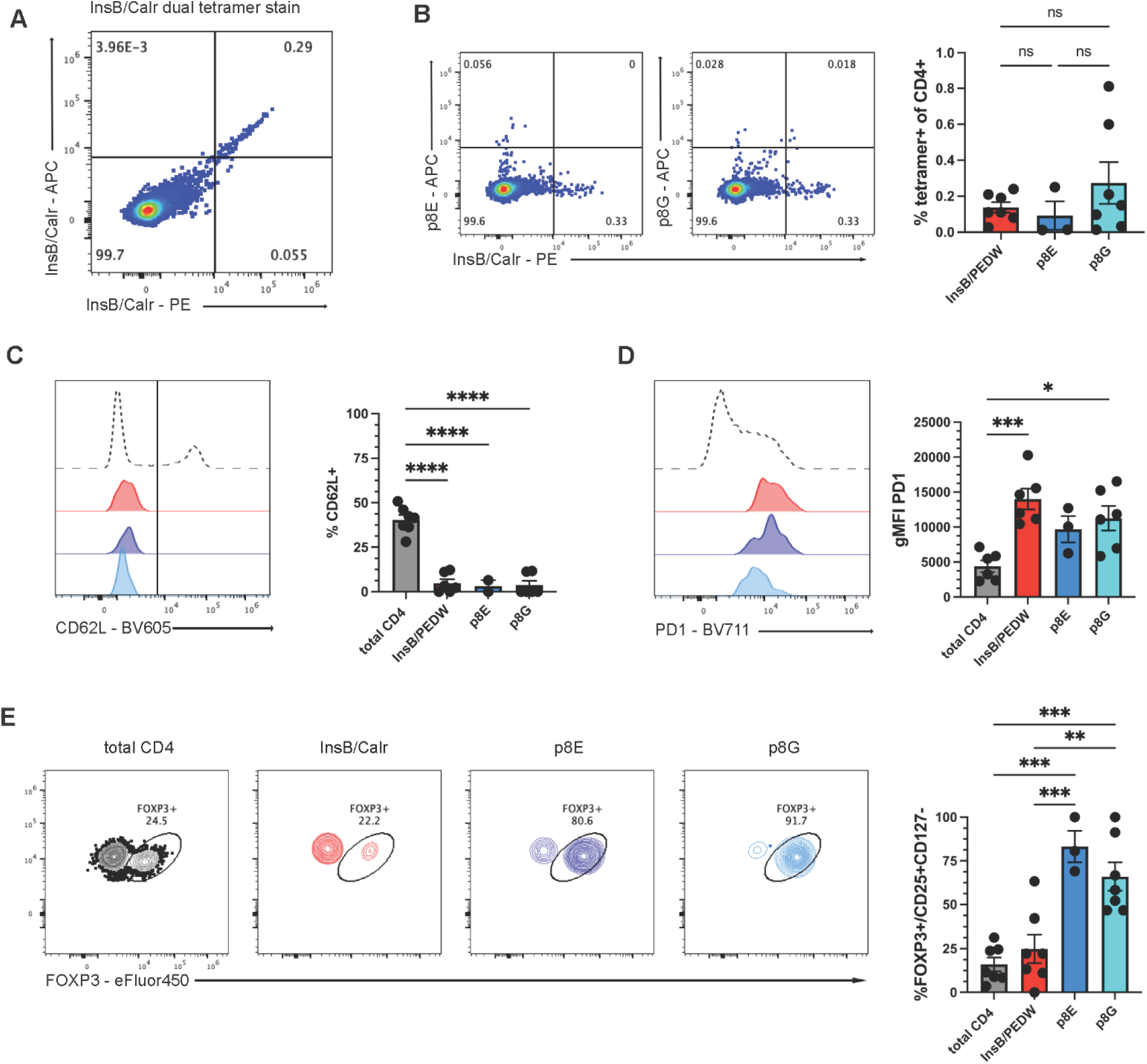
InsB/Calr-specific CD4 T cells are present within pancreatic islets of NOD mice. (A) Representative flow cytometry plots of dual tetramer staining of CD4 T cells isolated from NOD islets using PE- and APC-labeled I-A^g^^7^ tetramers loaded with the InsB/Calr peptide. (B) InsB/Calr tetramer staining compared to InsB p8E and p8E tetramer staining and the % tetramer positive of total islet infiltrating CD4 T cells. Statistical analysis by ordinary one-way ANOVA with Tukey’s multiple comparison test. (C) Expression of CD62L on tetramer+ (InsB/Calr, p8E, p8G) versus tetramer-CD4⁺ T cells isolated from NOD islets. (D) PD-1 gMFI expression on tetramer+ vs tetramer-CD4 T cells isolated from NOD islets. Data are pooled from two independent experiments using islets from 3 mice per experiment. (E) FOXP3 expression on CD4+ vs tetramer+ (insB/Calr HIP, p8E, p8G) isolated from NOD islets. In B-E, data are presented as the mean ± SEM, statistical analysis by ordinary one-way ANOVA with Tukey’s multiple comparison test, adjusted p-value <0.05 =*, <0.01 =**, <0.01 = ***, <0.001 = ***, <0.0001 = *****.

Phenotypic analysis demonstrated that InsB/Calr HIP tetramer+ cells exhibited an antigen-experienced phenotype characterized by an absence of CD62L expression compared to total CD4 T cells (**Fig. 4B**). InsB/Calr HIP-specific T cells also expressed elevated levels of PD1 (**Fig. 4C**), suggesting repeated antigen encounter within the inflamed islet environment. Only a small subset of InsB/Calr HIP tetramer+ cells expressed a regulatory phenotype (FOXP3+ or CD127-CD25+) compared to their p8E and p8G counterparts (**Fig. 4D**), suggesting that there are few regulatory CD4 T cells recognizing this B-chain HIP neoepitope within islets. These data demonstrate that InsB/Calr-specific CD4 T cells are present within inflamed NOD islets and exhibit an antigen-experienced phenotype, supporting this B-chain HIP as a physiologically relevant autoantigen.

### CD4 T cells from a recent-onset Type 1 Diabetic patient respond to the InsB/Calr HIP

To evaluate whether the InsB/Calr hybrid peptide identified in the NOD mouse model is also recognized by human T cells, we assessed antigen-specific proliferation in peripheral blood mononuclear cells (PBMCs) from recent-onset T1D donors (**Table 1**; human demographics). Because disease-associated HLA class II molecules such as HLA-DQ2 and HLA-DQ8 are known to present insulin-derived epitopes and share peptide-binding preferences with I-A^g^^7^, PBMCs from donors carrying these alleles were selected for analysis. Cells were labeled with CFSE and stimulated with the InsB/Calr peptide or native, control peptides *in vitro*. After seven days of culture, proliferation of CD4 T cells was assessed by activation (CD25^hi^) and CFSE dilution (**Fig. 5A** gating in **S3C**). CD4 T cells from one recent-onset T1D donor (patient 1187) exhibited proliferative responses to the InsB/Calr peptide that exceeded those observed with corresponding native insulin B-chain and/or Calreticulin sequences (**Fig. 5A, B**). Individual CD4 T cells were subsequently cloned from the PBMCs of patient 1187. Three CD4 T cell clones and a polyclonal CD4 line were isolated that were reactive to the InsB_12-19_/Calr_250-256_ HIP (**Fig. 5C**). To further examine T cell responses to the InsB_12-19_/Calr_250-256_ HIP, one of the CD4 T cell clones which reacted strongly to the InsB_12-19_/Calr_250-256_ HIP (B7 clone) was selected for further characterization. This T cell clone reacted to the InsB_12-19_/Calr_250-256_ HIP but not to native insulin peptides (insB_8-26_), insB_9-23_, insB_9-23R22E_ mimotope or to the native Calr_242-256_ peptide (**Fig. 5D**). These results demonstrated that a portion of human, B-chain HIP reactive T cells can exclusively react to HIPs and not the native “donor” peptides that compose the HIP. Our data indicate that the response to the InsB_12-19_/Calr_250-256_ HIP is restricted to HLA-DQ as T cell responses were blocked in the presence of the anti-DQ antibody SPV-L3 but not anti-DR antibody L243 (**Fig. 5E**). Further analysis indicates that the InsB_12-19_/Calr_250-256_ HIP can be presented in the context of HLA DQ2.5 (DQA1*0501 : DQB1*0201) as mis-matched EBV-transformed B cell lines only carrying the HLA DQ2.5 allele stimulated the B7 CD4 T cell clone in the presence of InsB_12-19_/Calr_250-256_ HIP and this activation was blocked with a pan anti-DQ antibody (**Fig. S6A**). Titration of the InsB/Calr HIP incubated with either autologous or DQ2.5-expressing APCs produced similar stimulation of the B7 clone (**Fig. S6B**), further indicating that the B7 TCR is DQ2.5-restricted. RNA sequencing of these clones (**Fig. S6C**) followed by TRUST4-based TCR imputing^42^ showed that clones B7, G11, and D2 all contain matched TCR alpha chains (TRAV21*01,TRAJ17*01, CDR3: CAVHKAAGNKLTF), with no generated data for TCR beta. Clone B4 showed distinct TCRalpha (TRAV9-2*01, TRAJ38*01, CDR3: CALINAGNNRKLIW) and TCRbeta (TRBV7-6*01, TRBD2*01, TRBJ2-1*01, CDR3: CASSPTGGYYNEQFF) chains, suggesting that multiple TCR clones respond to the InsB/Calr HIP. Together, these results obtained from analysis of T1D donor PBMCs indicate that the InsB/Calr HIP can be recognized by human CD4 T cells and support the relevance of insulin B-chain-derived HIP epitopes as targets of autoreactive T cells in T1D.

**Figure 5.**
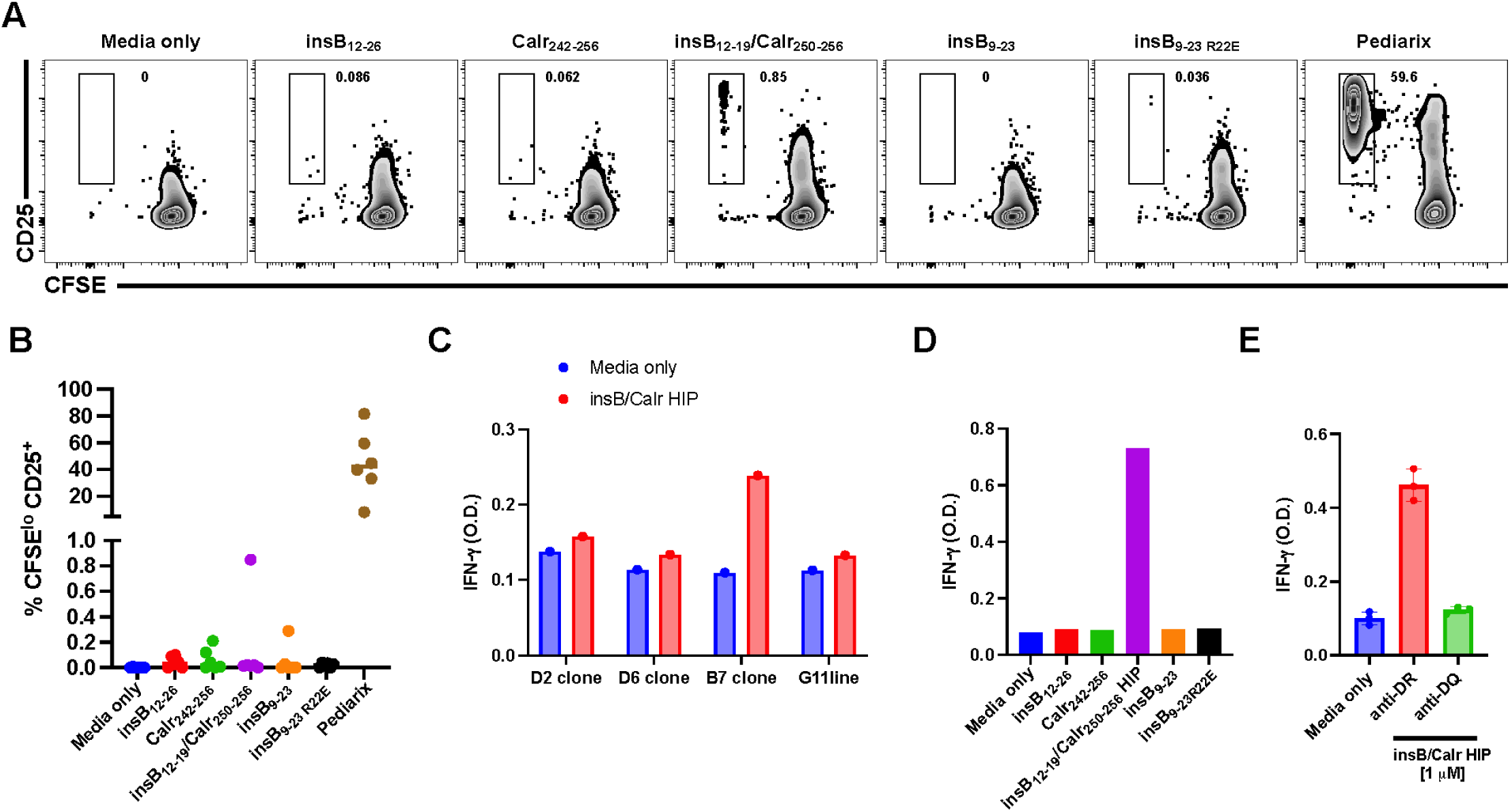
CD4 T cells from recent-onset T1D donors proliferated more robustly in response to the InsB/Calr hybrid peptide than to native insulin B-chain peptides. **(A)** Peripheral blood mononuclear cells (PBMCs) from recent-onset T1D donors (patient 1187) were labeled with CFSE and stimulated with native insulin B12–26, the native Calreticulin242–256, InsB/Calr HIP; (InsB_12-19_/Calr_250-256_), native InsB_9-23_, the InsB_9-23_ mimotope, or the recall vaccine Pediarix for 7 days. Representative flow cytometry plots show CFSE dilution within the CD4⁺ T-cell population following culture in the absence (media only) or presence of antigen stimulation. Cells were gated on live CD4⁺CD8⁻ lymphocytes within the FSC^int^/SSC^lo^ population. **(B)** Summary of proliferative responses from six recent-onset T1D donors carrying HLA-DQ2 and/or HLA-DQ8. Each point represents an individual donor. **(C-E)** IFN-γ ELISA results from culture supernatants collected after 48 h of stimulation under the indicated conditions using T-cell clones or T-cell lines isolated from patient 1187. **(C)** Ninety-nine individual T cells and one polyclonal T-cell line were sorted by FACS from patient 1187. Three T-cell clones and one polyclonal T-cell line were successfully expanded. The B7 CD4⁺ T-cell clone exhibited the strongest response and was selected for further characterization. Data are from a single experiment and are representative of 2 independent experiments. **(D)** IFN-γ responses of the B7 CD4⁺ T-cell clone to the indicated peptides. Data are from one experiment and representative of three independent experiments.**(E)** The B7 CD4⁺ T-cell clone was cultured in the absence of antigen (media only) or in the presence of antigen together with blocking antibodies specific for HLA-DR (anti-DR) or HLA-DQ (anti-DQ). Data are summarized from three independent experiments. Vertical bars represent the mean ± SEM.

**Table 1.** Human Demographic Table. Demographics of human patient data in figure 5 including patient study ID, HLA sequence, autoantibody positivity, sex, time from T1D diagnosis to analysis, age, ethnicity, and race designation.

## Discussion

Hybrid insulin peptides (HIPs) represent an emerging class of post-translationally generated neoantigens recognized by autoreactive T cells in T1D^21^. In this study, we identified several previously undescribed HIPs formed by the fusion of insulin B-chain sequences with fragments derived from other β-cell proteins. A specific HIP formed by fusion of insulin B-chain residues with a calreticulin-derived fragment was recognized by multiple TCRs cloned from islets of NOD mice. We then demonstrated that T cells recognizing this epitope are present within the islet infiltrate of NOD mice in an antigen-experienced, non-regulatory state. Lastly, we showed that autoreactive CD4 T cells recognizing this InsB/Calr B-chain HIP can be detected in a new onset T1D human patient. These findings demonstrate the utility of SABR screens to identify neoantigens in autoimmunity and expand the set of insulin-derived HIP epitopes, further supporting the concept that peptide fusion events that form neoepitopes contribute to B-chain antigen pool in autoimmune diabetes.

Our findings extend this paradigm by identifying calreticulin as a donor peptide for HIP formation. Calreticulin is an endoplasmic reticulum chaperone involved in protein folding and calcium homeostasis^43^, and fragments derived from this protein may enter antigen processing pathways during β-cell stress or inflammation. The identification of a calreticulin-derived HIP, the sequence of which is identical between mice and humans, suggests that a broader range of intracellular proteins may contribute to HIP formation than has been previously appreciated. Importantly, residues contributed by the calreticulin segment played a critical role in determining antigenicity of the peptide. Substitution of acidic residues predicted to occupy the p8 and p9 positions of the peptide-MHC complex significantly reduced TCR activation, indicating that the donor peptide contributes residues that potentially stabilize peptide-MHC binding and enhance TCR binding. In this way, HIP formation may enhance the ability of insulin-derived fragments to bind disease-relevant MHC molecules. Because insulin B-chain peptides can bind MHC class II molecules such as I-A^g^^7^ (or HLA-DQ8) in multiple registers, fusion with donor peptides may introduce residues that stabilize particular binding registers by improved anchoring within the MHC binding groove. Thus, hybrid peptide formation may endow insulin B-chain fragments with increased capacity to engage disease-associated MHC molecules in a prolonged manner and thus activate autoreactive T cells within islets.

Consistent with this interpretation, InsB/Calr HIP-specific CD4 T cells were detected within NOD islets and displayed an antigen-experienced phenotype characterized by elevated PD1 expression and reduced CD62L, indicative of repeated antigen encounter in the inflamed islet environment. The frequency of InsB/Calr HIP tetramer-positive cells was comparable to that observed for other insulin B-chain-reactive tetramer populations, suggesting that InsB/Calr HIP epitope represents a substantial component of the autoreactive CD4 T cells, similar to our previous findings with another B-chain-derived HIP^30^. Interestingly, only a small fraction of InsB/Calr tetramer+ cells were FOXP3+/CD127-CD25+ compared to insulin B-chain p8G and p8E tetramer+ cells. This suggests that regulatory T cell responses to this epitope may be relatively limited and pathogenic effector T cells are predominant, similar to what has been observed for 2.5HIP-reactive CD4 T cells^23^. The presence of T cells that either react only with InsB/Calr HIP, or cross-react with InsB/Calr HIP and InsB:9-23, suggests that the neoantigenicity of this HIP may occur through two potential mechanisms. One possibility is that the HIP acts as a superagonist to CD4 T cells weakly reactive to insulin B-chain that escaped negative selection. Another possibility is through stimulation of de novo peripheral repertoires that would not be subjected to central tolerance, similar to 2.5HIP-reactive T cells. Further studies comparing the TCR repertoires and phenotypes of these cells are warranted to distinguish between these possibilities.

Our study also provides evidence that the InsB/Calr HIP is recognized by human T cells. CD4 T cells from a recent-onset T1D donors carrying HLA-DQ2 proliferated in response to the insB/Calr HIP in CFSE-based assays, exceeding responses to native insulin B-chain or Calreticulin peptides. One intriguing finding was that this response was restricted to HLA-DQ2. In a previous study, we have found that HIP11, a hybrid insulin peptide formed between two C-peptide fragments is also presented by HLA-DQ2, highlighting the importance of this high-risk HLA molecule in the presentation of neoepitopes. Determining the kinetics of these HIP-specific T cell responses before and during disease progression will be important to understand how they arise relative to other characterized β-cell autoantigens. Together, these findings suggest that hybrid peptides derived from the insulin B-chain may be targets of autoimmune T cell responses in both mice and humans.

More broadly, these findings highlight the power of SABR-based screening platforms for identifying antigenic ligands of autoreactive T cells. Traditional peptide screening approaches are limited by throughput and cannot efficiently interrogate the vast combinatorial space of potential hybrid peptides or other PTMs. By contrast, SABR-based libraries enable functional screening of thousands of candidate peptides simultaneously, providing a powerful strategy for discovering previously unrecognized autoantigens. Identifying the full panoply of native and PTM epitopes recognized by autoreactive T cells in T1D may provide sensitive biomarkers of disease kinetics and inform the development of antigen-specific immunotherapies aimed at restoring tolerance to β-cell antigens.

Demographics of human patient data in figure 5 including patient study ID, HLA sequence, autoantibody positivity, sex, time from T1D diagnosis to analysis, age, ethnicity, and race designation.

## Methods

### Mice

Female NOD/ShiLtJ mice (cat. No. 001976) were obtained from The Jackson Laboratory and bred inhouse for no more than 5 generations before new breeders were reordered. Animals were maintained under specific pathogen-free conditions (12/12 light/dark cycle, 40-60% humidity, food and water ad libitum) either in the animal facilities at the University of Pittsburgh or at the University of Colorado Anschutz Medical Campus. Mice were used between 8 and 14 weeks of age, corresponding to the prediabetic stage when insulitis is established but overt diabetes has not yet developed. All animal experiments were conducted in accordance with institutional guidelines and were approved by the Institutional Animal Care and Use Committee (IACUC) at University of Pittsburgh and University of Colorado Anschutz Medical Campus.

### Isolation of pancreatic islets and preparation of islet-infiltrating lymphocytes

Single cell RNA and TCR sequencing data were obtained as previously described^33^. For flow cytometry experiments, pancreata were inflated through the common bile duct with CIzyme RI (VitaCyte), dissolved in HBSS, and incubated at 37°C for 17 min. Digested tissue was washed in cold HBSS and islets were purified using a lympholyte (Cedarlane labs) density gradient. Islets were then washed and dispersed with non-enzymatic cell stripper (Thermo) for 10 minutes with intermittent mixing by pipetting at 37°C. Cells were then washed in 10% FBS RPMI prior to staining.

### Design and construction of hybrid insulin peptide library

A theoretical HIP library was constructed by combinatorial pairing of insulin B-chain fragments with peptides derived from β-cell secretory proteins. The N-terminal portion of the library consisted of seven-amino acid truncations spanning insulin B-chain residues 6-20 derived from Ins1 and Ins2 sequences. The C-terminal portion consisted of 6-8 amino acid fragments derived from β-cell proteins known to localize to insulin secretory granules including insulin, chromogranin A, secretogranin-1, secretogranin-2, IA-2, Znt8, calreticulin, and others (full list of left and right peptides in supplemental data). The resulting library consisted of 7,884 unique hybrid peptide sequences and 3 additional controls. Oligonucleotides encoding for the peptide sequences were flanked with overhangs and cloned into the SABR backbone. The library was synthesized by Twist Biosciences using pooled synthesis. Oligonucleotide pools were amplified by PCR using a protocol previously described^34^, and cloned using Infusion Cloning HD (Takara Bio) into BsmBI-digested (NEB) I-A^g^^7^ SABR backbone. HIP libraries were verified for >100x coverage by Sanger sequencing, and expressed in NFAT-GFP-Jurkat cells. SABR expression was confirmed by flow cytometry (Anti-RT1B-PE, Biolegend).

### SABR screening

SABR libraries were screened as previously described ^33,34^. SABR-expressing cells were stained with Celltrace Violet (Life Technologies), and co-incubated with Jurkat cells expressing individual insulin-reactive TCRs. Following co-culture, cells were stained with anti-CD69 APC-Cy7 (Biolegend), and GFP+CD69+ cells were isolated by fluorescence-activated cell sorting (FACS) on BD Aria III (BD Biosciences). Genomic DNA was extracted from sorted cells using PureLink genomic DNA kit (Life Technologies). The SABR portion of the epitope was amplified, indexed, and subjected to next-generation sequencing to identify enriched sequences. Enrichment scores were calculated as previously described, using a linear model of expected read fractions for each epitope across sorted cells in each sample vs the unsorted library. A stringent cutoff for enrichment was used to identify putative epitopes. Candidate peptides identified in the screen were cloned individually into SABR constructs using oligonucleotides as per previous protocols ^33,34^ and tested for their ability to activate TCR reporter cells.

### Reporter cell activation assays

Jurkat-based cell lines expressing TCR11, TCR24, or TCR30 were used to assess activation by candidate hybrid peptides SABRs cells. TCR expressing-cells were co-cultured with NFAT/GFP-reporter SABR-expressing cells presenting individual hybrid insulin peptides. After 18-24 h of co-culture, reporter activation was assessed by flow cytometry measuring GFP reporter expression and CD69 surface expression. Cells were analyzed using Attune NxT flow cytometer (Life Technologies) and data were analyzed using FlowJo v10 software. For peptide mapping experiments, truncation and substitution variants of the InsB/Calr peptide were cloned into SABR constructs and tested in the same assay.

### Tetramer staining

Peptide-MHC class II tetramers were generated by loading recombinant I-A^g^^7^ molecules with the InsB/Calr peptide (VEALYLVC/PEDWDEE) or the InsB/Scg1 peptide (VEALYLVC/SEEEPEY) at the NIH tetramer core. Peptides were synthesized at >95% purity from GenScript (NJ, USA) and used for tetramer generation. Tetramers were conjugated to PE or APC fluorophores. Islet-derived lymphocytes were stained with tetramers 20 minutes then surface antibodies were added in for an additional 40 min at 37°C 5% CO_2_. Cells were washed and stained for viability, and either analyzed by flow cytometry or fixed for intracellular staining. Intracellular FOXP3 staining was performed using the Foxp3/Transcription Factor Staining Buffer Set (eBioscience) per the manufacturer’s recommendations. Cells were analyzed by flow cytometry using a Cytek Aurora 5-laser instrument.

### Human PBMC proliferation assays and cloning

Peripheral blood mononuclear cells (PBMCs) were obtained from recent-onset T1D donors carrying HLA-DQ2 or HLA-DQ8 alleles through the University of Colorado clinical research program (IRB 01-384). PBMCs were labeled with CFSE (Thermo Fisher Scientific) according to the manufacturer’s instructions. Labeled cells were cultured in AIM V supplemented with 2.5% human AB serum (Gibco) and stimulated with InsB/Calr HIP (10 μg/ml), control peptides or Pediarix®. After seven days of culture, cells were stained with antibodies against CD8 (BB700), CD4 (PE or BV711), CD45RA (BV510), CD45RO (PE-Cy7), CD25 (BV421) and a Fixable Viability Dye (FVD ef780) and analyzed by flow cytometry (Fortessa X20, BD Biosciences, USA). Proliferation and activation of CD4 T cells was assessed based on CFSE dilution and CD25 upregulation. For T cell cloning, methods were adapted from previously published protocols. Briefly, PBMCs were labeled with CFSE and cultured for 7-8 days in AIM V containing InsB/Calr HIP (10 μg/ml) and 2.5% normal human AB serum. CD4^+^ CD8^−^ CFSE^dim^, CD25^hi^, and ef780^neg^ cells were index sorted into a round-bottom 96-well plate using a FACS Aria cell sorter (BD Biosciences, USA) at one cell per well or 201 cells when generating polyclonal T cell lines. Wells were supplemented with recombinant human IL-2 (20 units/mL), IL-4 (5 ng/mL), anti-CD3 (30 ng/mL), irradiated mixed-matched PBMCs from two different donors (1 X 10^6^/mL), and irradiated PRIESS cells (1 X 10^6^/mL). Wells were regularly scored for growth by visual inspection, and cells were expanded using IL-2 and IL-4. After 28 days, T-cell clones were challenged with peptides using autologous irradiated PBMCs, and supernatants were tested for IFN-γ by ELISA. Wells that gave a positive response with little background were selected for further expansion and T-cell receptor (TCR) sequencing. For TCR sequencing, cell pellets from the individual clones were lysed in Zymo DNA/RNA Shield Stabilization solution and subjected to RNA seq (Plasmidsauraus). Resulting Fastq files were processed with TRUST4 to impute TCRalpha/beta chains.

### Flow cytometry

All flow cytometry experiments were performed using a Aurora 5-laser instrument (Cytek), a Fortessa X20 (BD Biosciences), or a 4-laser Attune NxT flow cytometer. Data were analyzed using FlowJo software v10 (Tree Star) and gating strategies for individual experiments can be found in Figure S3.

### Statistical analysis

Statistical analyses were performed using GraphPad Prism software. Data are presented as mean ± SEM unless otherwise indicated. Comparisons between multiple groups were performed using a one-way ANOVA with post Tukey’s multiple comparison correction as appropriate. For human PBMC proliferation assays, nonparametric tests were used when sample sizes were small. A p-value <0.05 was considered statistically significant. Islet samples were excluded from analysis if they had too few total CD4 T cells (<5,000 cells/islet samples).

## Author contributions

Designing research studies (JED, PMZ, JMW, KH, RLB, AVJ), conducting experiments (JED PMZ, NT, JMW, ACH, JG, RLB), acquiring data (JED, PMZ, NT, JMW, ACH, JG, RLB), analyzing data (JED, PMZ, NT, RLB, AVJ), providing reagents (JG, PAG, KH, RLB, AVJ), writing the manuscript (JED, KH, RLB, AVJ).

JED and PMZ share co-first authorship and contributed equally to the experiments, JED is listed first because of their contribution to the final writing and figures within the manuscript.

## Acknowledgments

We would like to acknowledge the Barbara Davis Center Tissue Procurement and Processing Core at the University of Colorado Diabetes Research Center, NIDDK Grant P30-DK116073. We would like to acknowledge the Cancer Center Support Grant (P30CA046934) and the Flow Cytometry Shared Resource at the University of Colorado Anschutz Medical Campus. We are grateful for the open-source images available from the NIAID NIH BioArt Source. We are thankful for the reagents provided by the NIH Tetramer Core Facility (NIH Contract 75N93020D00005 and RRID:SCR_026557). We would like to acknowledge the University of Pittsburgh Unified Flow Core and Division of Laboratory Animal Research staff. We thank L. Kublo and E. Zarate Martinez for technical help with molecular biology and Kelli Nicholson for help with T cell cultures.

## Additional reagent information

List of peptides, antibodies, and reagents can be found in **Table S1**.

