## Supplementary material for "Cell-based epitope screen identifies a novel B chain-derived Hybrid Insulin Peptide recognized by human and mouse autoreactive CD4 T cells"

### Supplemental Figures

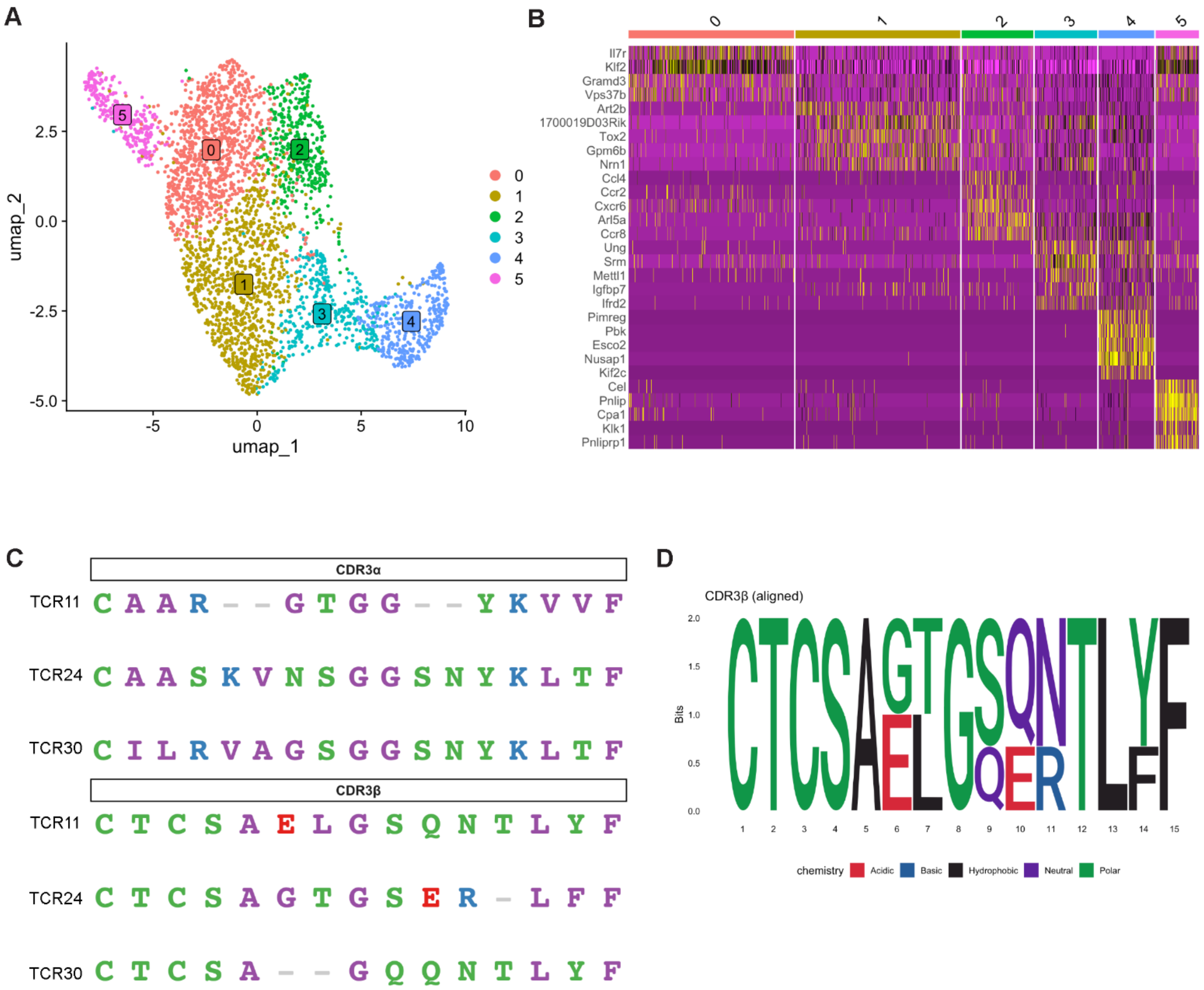


**Figure S1. scRNAseq cluster defining genes and CDR3 alignment.**
(A) UMAP cluster of CD4 T cells from NOD mouse islets.
(B) Cluster defining genes.
(C) TCR CDR3α and CDR3β amino acid alignment from TCR11, TCR24, and TCR30.

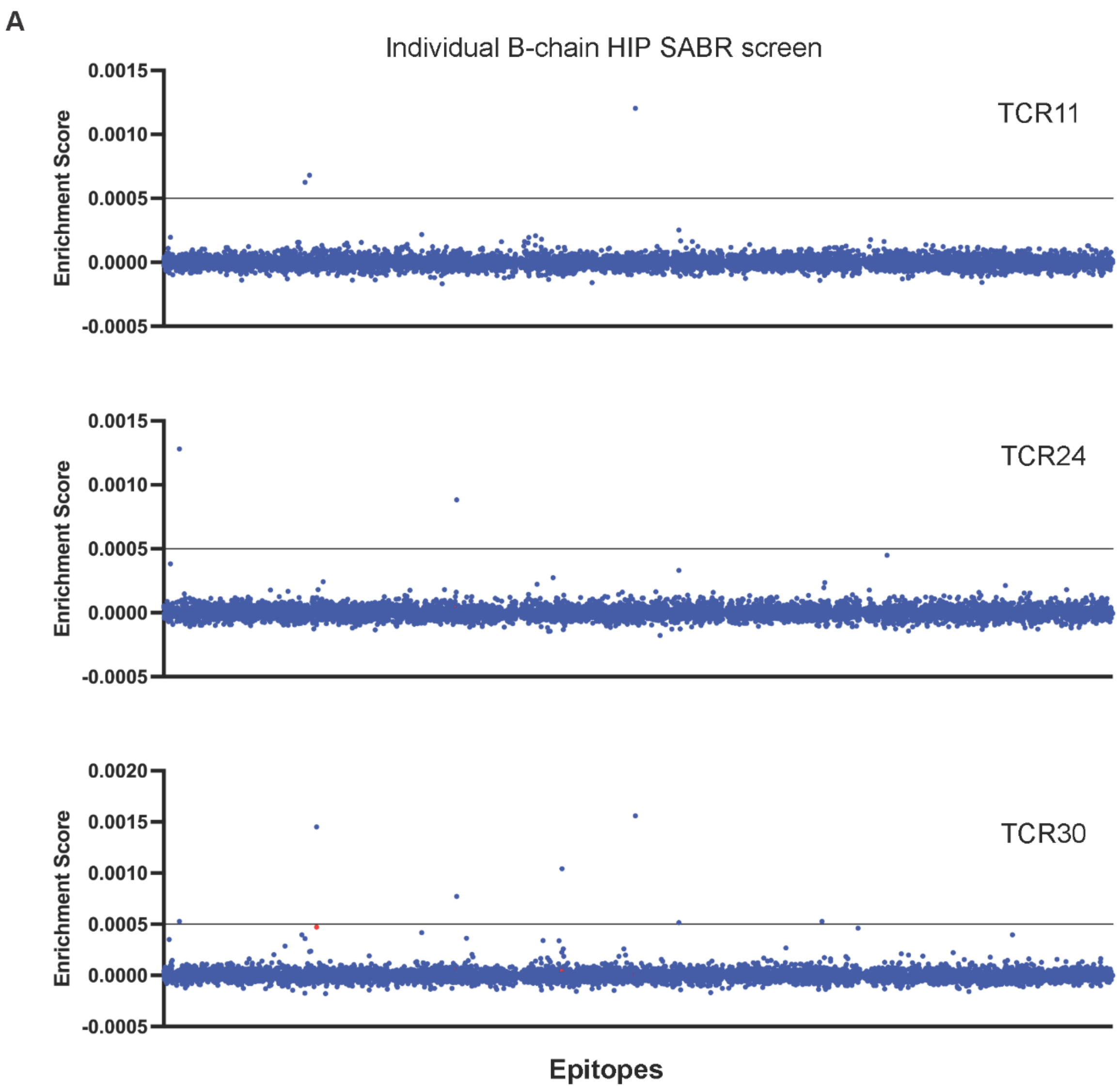


**Figure S2. Individual SABR screen each TCR alone TCR11, TCR24, TCR30.**
(A) Enrichment plots for each individual TCR on the B-chain HIP SABR library.

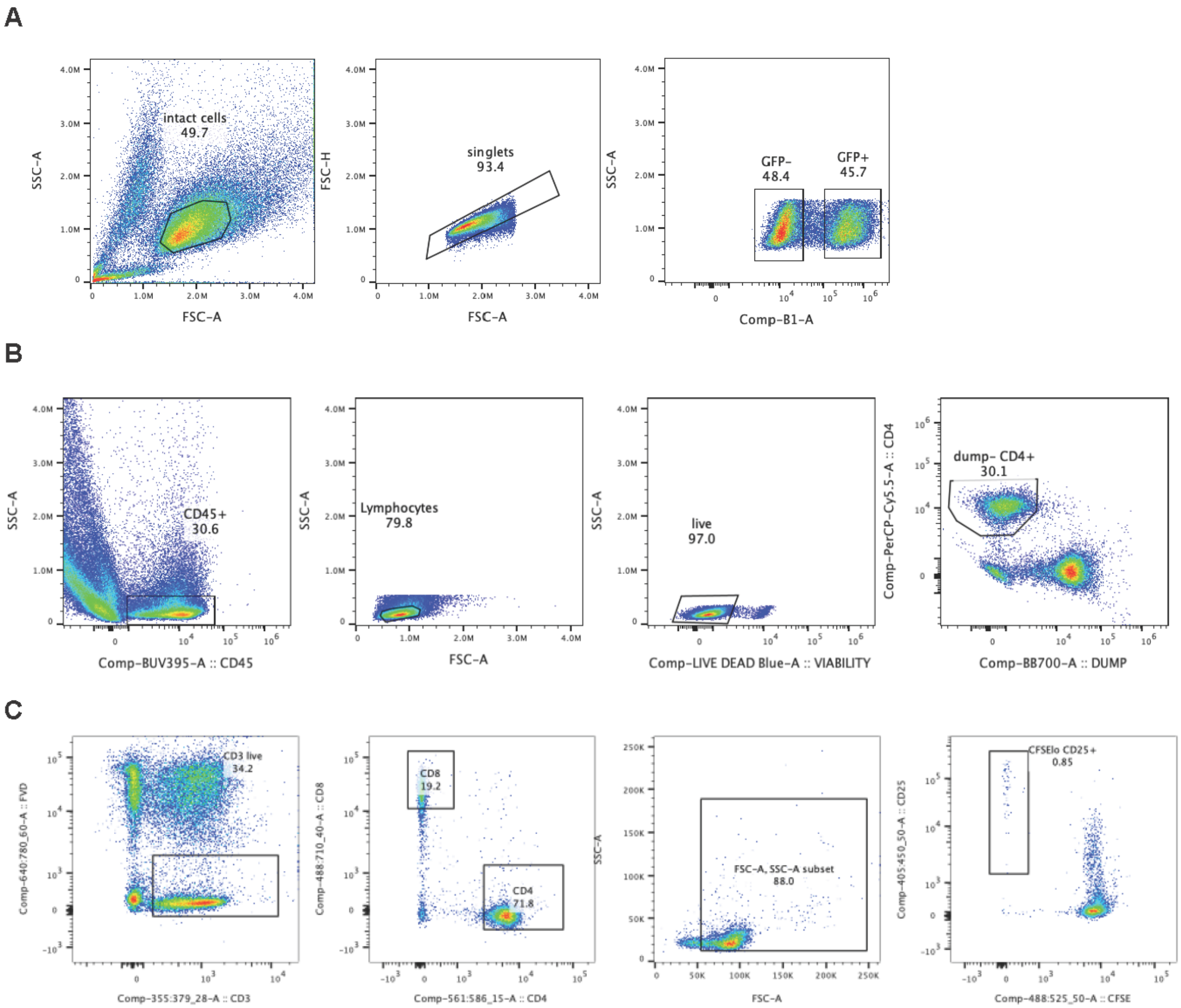


**Figure S3. Flow cytometry gating strategies**

(A) Gating strategy for Jurket cells expressing individual TCR constructs (Figure 2E). Intact cells, singlets, GFP- (no TCR construct), GFP+ (with TCR construct).

(B) Gating strategy for islet infiltrating CD4 T cells (Figure 4A-E). CD45+, lymphocytes, live, lineage- (CD11c, CD11b, Ly6G, CD19, CD8a) and CD4+.

(C) Gating strategy for human PBMC CD4 T cells. Gates were set on Live (FVD$^-$) T cells (CD3$^+$) lymphocyte (FSC$^{int}$; SSC$^{lo}$), CD8$^-$ , CD4$^+$ cells. The percentages of CFSE$^{lo}$ CD25+ cells are reported. (Figure 5).

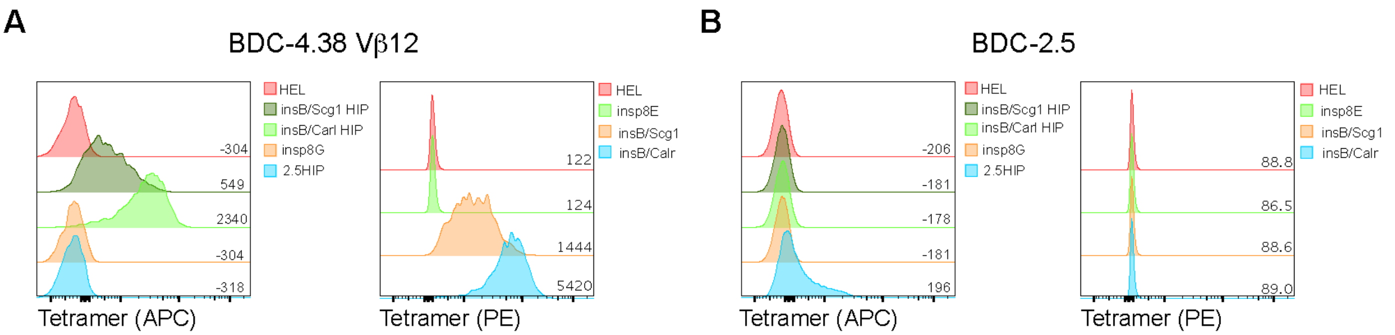


**Fig. S4. The BDC-4.38 Vb12 CD4 T cell clone does not bind the insp8G nor insp8E tetramers.** (A) The BDC-4.38 Vb12 T cell clone was stained with CD4 BB700, the Fixable Viability dye ef780 and individual insulin tetramers. Data are from one experiment and are representative of 2 independent experiments.

A

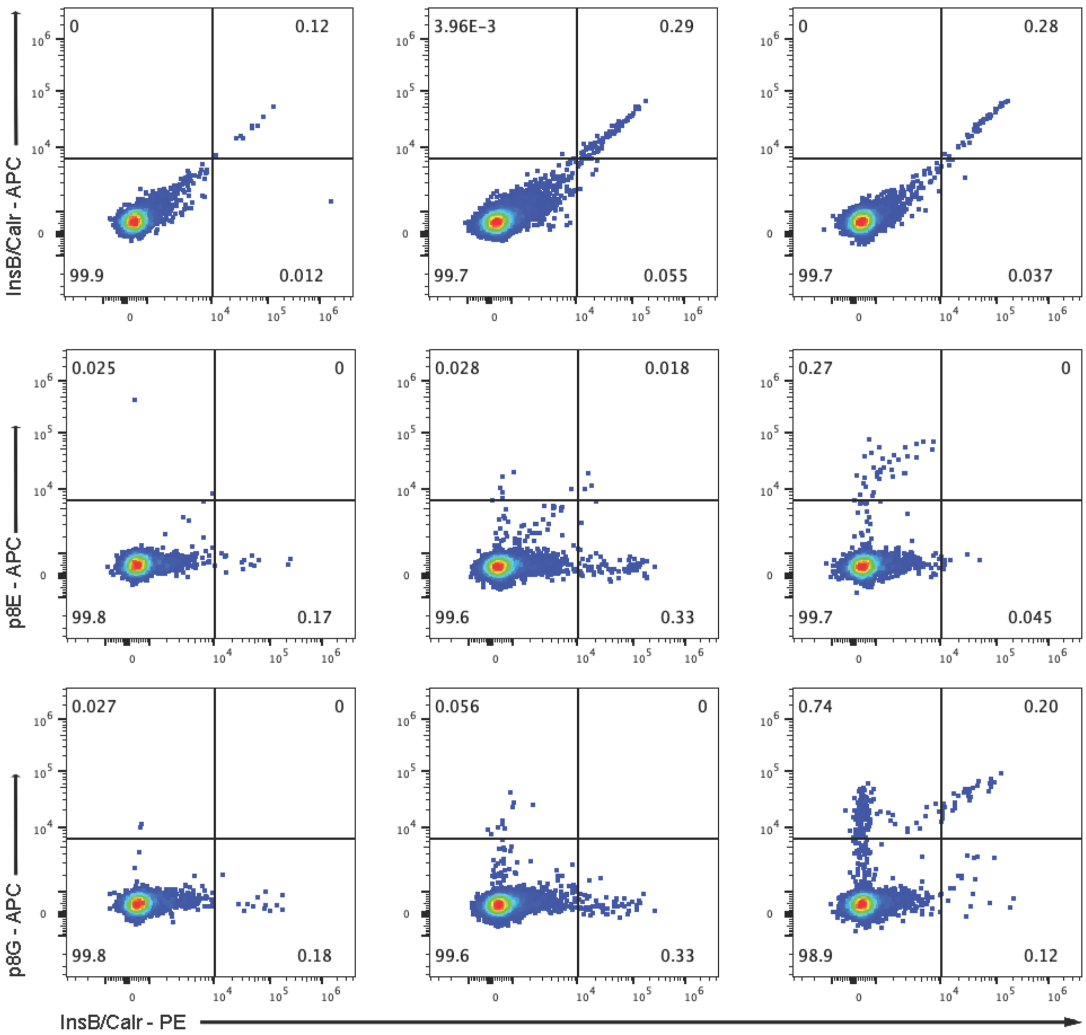


**Figure S5. InsB/Calr, p8G, and p8E tetramer stains of three individual animals**

(A) Flow plots of CD4 T cells pre-gated as in Figure S3A from the islets of three individual animals co-stained with the InsB/Calr HIP tetramer in PE by either the InsB/Calr HIP, p8E, or p8G tetramers in APC.

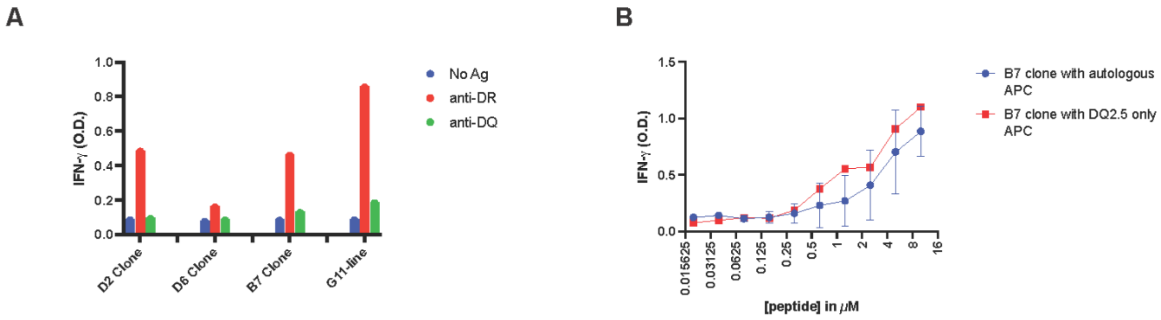


C

| | | | | |
|---|---|---|---|---|
| Clone B4 | CASSPTGGYYNEQFF | TRBV7-6*01 | TRBD2*01 | TRBJ2-1*01 |
| | CALINAGNNRKLIW | TRAV9-2*01 | | TRAJ38*01 |
| Clone B7, G11, D2 | CAVHKAAGNKLTF | TRAV21*01 | | TRAJ17*01 |

**Figure S6. HLA restriction of InsB/Calr HIP-reactive human T cell lines and clones**

(A) T cells from patient BDC1187 were incubated with EBV-transformed B cell lines with the HLA DQ2.5 allele and incubated with InsB$_{12-19}$/Calr$_{250-256}$ HIP in the presence of blocking antibodies specific for pan-human DR or DQ HLA haplotypes. Following peptide incubated, supernatants were collected and an IFN$\gamma$ ELISA was performed to measure T cell activation.

(B) The B7 CD4 T cell clone was incubated with either autologous EBV-transformed B cells or EBV-transformed B cells partially matched for HLA-DQ2 in the presence of indicated concentrations of the insB/Calr HIP. Levels of IFN-$\gamma$ was measured 48h IFN-$\gamma$ after co-culture. Data are a summary of two independent experiments. Experiment with partially matched EBV-transformed line was performed three times at the highest concentration (10 $\mu$M); titration experiment was performed once.

(C) CDR3 sequences following TCR sequencing of clones B4, B7, G11, and D2.

| Peptide | Name | Sequence | Company |
|---|---|---|---|
| insB12-26 | insB12-26 | VEALYLVCGERGFFY | GenScript |
| Calr242-256 | Calr242-256 | IPDPDAKKPEDWDEE | GenScript |
| insB12-19/Calr250-256 HIP | insB/Calr HIP | VEALYLVCPEDWDEE | GenScript |
| insB9-23 | insB9-23 | SHLVEALYLVCGERG | GenScript |
| insB9-23 R22E | insB9-23 R22E | SHLVEALYLVCGEEG | GenScript |

| Antibody | catatog # | Company |
|---|---|---|
| BV421 Mouse Anti-Human CD25 | 562442 | BD Horizon |
| PE-Cy7 Mouse Anti-Human CD45RO | 560608 | BD Pharmingen |
| BV510 Mouse Anti-Human CD45RA | 563031 | BD Horizon |
| BV711 Mouse Anti-Human CD4 | 563028 | BD Horizon |
| BB700 Mouse Anti-Human CD8 | 566452 | BD Horizon |
| PE Mouse Anti-Human CD4 | 561843 | BD Pharmingen |
| BUV395 Mouse anti-Human CD3 | 563546 | BD Horizon |
| Fixable Viability Dye ef780 | 65-0865-14 | ThermoFisher Scientific |

| Reagent | | Company |
|---|---|---|
| AB serum | 100-512 | Gemini |
| AIM V | 12055-091 | Gibco |

**Table S1 Additional reagent information**

(A) Table including peptides sequences, human antibody catalog numbers, and human culture reagents.
